# Emergence of rigidity percolation and critical behavior in tunable protein condensates

**DOI:** 10.64898/2026.04.13.718064

**Authors:** Zhitao Liao, Bowen Jia, Yang Xu, Zeyu Shen, Mingjie Zhang, Penger Tong

## Abstract

Multivalent proteins are known to form complex networks within biomolecular condensates, yet the mechanisms governing the emergence and evolution of these networks remain poorly understood. Here, we utilize a synthetic protein chimera system with tunable interactions to investigate the transition from liquid-like droplets to networked condensates. By employing single-amino-acid substitutions, we generated a series of mutants with varying protein-binding strengths. As the interaction strength increases, the condensates undergo a sharp rigidity percolation transition, characterized by a more than 200-fold increase in both elastic modulus and viscosity. Near this transition, we identify a critical scaling relation in the condensate elasticity, providing robust evidence for a percolation-driven assembly mechanism. Furthermore, we demonstrate that this network architecture is fundamentally linked to biological functions. The introduction of disease-associated mutations disrupts the network, significantly softening the condensates and rendering them fluid-like. Our findings reveal that network percolation and criticality in protein condensates can be sensitively regulated by single-amino-acid substitutions, underscoring their essential role in maintaining structural integrity and supporting physiological function.

## Introduction

Biomolecular condensates are phase-separated membraneless compartments in living cells that concentrate specific biomolecules [1–3]. Due to the complexity of molecular interactions, these condensates behave as complex, viscoelastic materials and exhibit diverse material properties across multiple spatiotemporal scales [4–6]. The material properties of the condensates, such as viscosity *η*, elastic modulus *E* and interfacial tension *γ*, are crucial in determining the condensate behavior and regulating biological activities [5–8].

When protein condensates contain multiple components or involve strong multivalent interactions, the condensed molecules can form network structures, giving rise to emergent material properties beyond simple Maxwell fluids [9–12]. These network-related properties are strongly correlated with physiological functions and homeostasis of the cells, like neuronal signal transmission [13, 14], and with condensate pathology such as aging-related diseases [15, 16]. For example, synaptic condensates feature soft-glass-like properties, which are critical to maintain the integrity and normal function of the synapse [17–19]. Weakening of this network can hinder its ability to anchor synaptic receptors and induce pathological changes in mice’s brain [19]. Abnormal protein aggregation during aging can generate pathological fibrillar networks and tangles, which are associated with various neurodegenerative diseases [15, 16, 20].

How does the network structure emerge from a condensation of proteins? A common network formation mechanism is through percolation [10, 11, 21–23]. Network percolation is a phase transition process initially introduced as a mathematical model of connectivity in graph or lat-tice, which refers to the emergence of a connected path when the bond connectivity (or site occupation) reaches a certain threshold [22–24]. Phase transitions are typically accompanied by abrupt changes in the macroscopic properties of the system when an external control parameter *p* reaches the threshold *p*_*c*_ [23, 25]. Near the transition, the system exhibit universal features characterized by power-law scaling as, *A* ∼ (*p− p*_*c*_)^*β*^, where *A* is an observable (such as magnetization or modulus), *p* is the control parameter (such as temperature or concentration), and *β* is the critical exponent [23, 25].

Recently, percolation theory has been widely adopted to describe abrupt mechanical transitions in various biological processes [10, 11, 26, 27], including tissue morphogenesis in zebrafish embryos [28] and the activation of complement proteins during immune responses [29]. Despite its growing application, targeted experimental studies that carefully examine the critical behavior of protein networks and quantify their critical exponents remain scarce. Whether biomolecular condensates exhibit sharp transitions in material properties and obey critical scaling laws during network formation is still an open question. While percolation is increasingly invoked to explain emergent phenomena in condensates, the specific mechanisms driving the formation of these network structures, and how they behave near the transition, have yet to be thoroughly elucidated.

In this study, we employ a synthetic protein chimera, designated PrLD-SAM, to investigate network percolation and the critical behavior of protein condensates. The PrLD-SAM system comprises a prion-like domain (PrLD) derived from the fused in sarcoma (FUS) protein and a folded sterile alpha motif (SAM) from the synaptic protein Shank3 [21]. The PrLD is an intrinsically disordered region (IDR) that mediates weak multivalent interactions [30, 31]; its role in regulating FUS phase separation is essential for maintaining nucleic-acid-related cellular functions [32, 33]. Conversely, the SAM is a folded domain that engages in specific head-to-tail interactions, forming polymeric chains that further assemble into bundles and sheets via side-chain interactions [34, 35]. This SAM-driven polymerization facilitates the construction of the postsynaptic density (PSD), a specialized synaptic condensate [13, 19, 36].

Consequently, SAM dysfunctions that impair polymerization can induce disease-associated synaptic changes, ultimately leading to various neurological disorders [37– 42]. We selected the SAM domain for this study because its interaction strength and polymerization capacity can be finely tuned via single-amino-acid mutations, allowing us to systematically examine how condensate mechanics evolves with protein interactions and to observe network percolation [19, 21, 35]. Thus, the PrLD-SAM chimera serves as a robust model system for mimicking functional proteins that utilize both IDR-mediated and folded-domain-induced multivalent interactions [21].

In this work, we show that the material properties of PrLD-SAM condensates undergo a rigidity percolation transition as SAM-SAM interactions are enhanced. This transition, evidenced by a dramatic increase in viscoelasticity, results from the emergence of a percolated network that forms only when the SAM interaction strength exceeds a critical threshold. Below this threshold, the condensates behave as Maxwell-like fluids, with their mechanics governed primarily by surface tension and viscosity. Above the transition, a system-spanning network emerges, providing the condensates with substantial elasticity against deformation. We also observe a power-law scaling in the condensate modulus, *E* ∼ (*ϕ− ϕ*_*c*_)^*β*^, which indicates clear critical behavior at the transition point. Furthermore, we find that network percolation is closely linked to physiological functions. Specifically, pathological mutations associated with neurological disorders can impair the network architecture, rendering the condensates fluid-like. Our findings not only demonstrate the emergence of network percolation and critical behavior in protein condensates but also highlight the crucial role of this network in maintaining structural integrity and regulating biological functions.

## Results

### Contrasting material properties between PrLD and wild-type SAM (SAMWT) condensates

Before discussing the material properties of the PrLD-SAMWT chimera condensate, we first examine the material properties of the condensates made by PrLD alone and by SAMWT alone. The PrLD and SAMWT proteins are end-capped by a pair of solubility tags, which are separated by an HRV-3C cut site, to enhance their solubility in an aqueous buffer before phase-separation (Fig. 1**a**, top panel). Upon cleavage of the two end-caps using HRV-3C protease, the two proteins phase separate into condensates. The PrLD forms spherical droplets, whereas SAMWT aggregates into fiber-like structures with irregular shapes or crystals (Fig. 1**a**, bottom panel). Fluorescence recovery after photobleaching (FRAP) was then used to measure protein mobility in the condensates. As shown in Figs. 1**b** and 1**c**, PrLD condensate exhibits rapid fluorescence recovery to near 100% within 2 minutes, suggesting PrLD molecules have a large diffusion coefficient *D*. By fitting the FRAP curve to a diffusion-driven model [43, 44], we find its characteristic recovery time *τ*_*r*_ = 11.7 s and *D* = 0.71 *µ*m^2^/s (see Supplementary Information (SI) Section I.D for details).

**FIG. 1.**
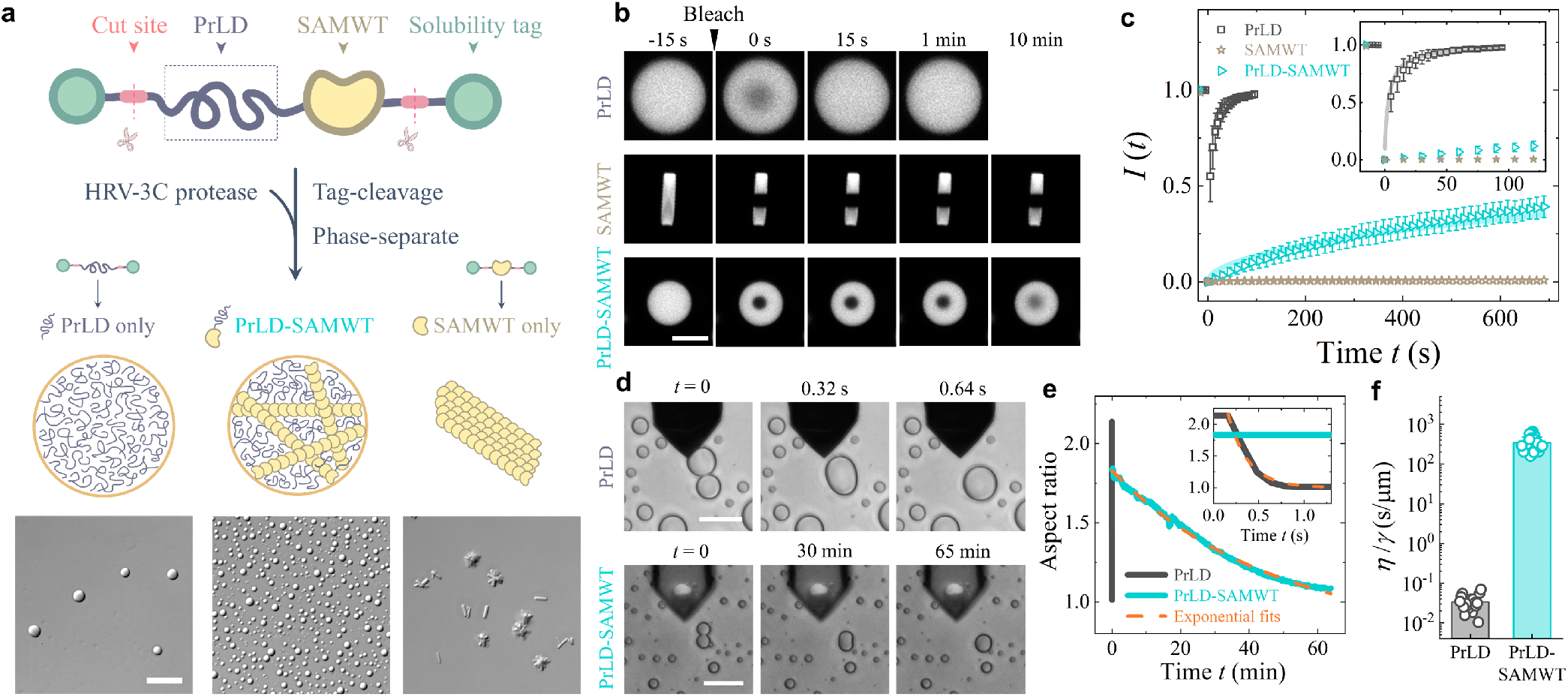
Material properties of PrLD, SAMWT, and PrLD-SAMWT condensates. **a**, Molecular design of PrLD-SAMWT chimera system. The PrLD, SAMWT, or PrLD-SAMWT molecules are end-capped by two solubility tags separated by an HRV-3C cut site (top panel). Upon cleavage of the two end-tags by adding HRV-3C protease in the buffer, the proteins phase separate into condensates (middle panel). The bottom panel shows the differential interference contrast images of the PrLD, PrLD-SAMWT, and SAMWT condensates. Scale bar: 20 *µ*m. **b**, Time-lapse confocal images of FRAP for the PrLD, SAMWT, and PrLD-SAMWT condensates. Scale bar: 5 *µ*m. **c**, Normalized fluorescence intensity *I*(*t*) from FRAP as a function of time *t* before (*t* < 0) and after (*t* > 0) photo-bleaching. The data (mean ± standard deviation) are obtained from *N* = 6 PrLD droplets, 9 PrLD-SAMWT droplets, and 8 SAMWT fibers. Inset shows a magnified view of the FRAP data at short times. **d**, Time-lapse images of PrLD and PrLD-SAMWT droplets showing the merging of two droplets into a single spherical droplet. The dark finger is the AFM cantilever. Scale bars: 20 *µ*m. **e**, Time evolution of the aspect ratio of merging droplets in **d** for the PrLD and PrLD-SAMWT condensates. Inset shows a magnified view of the aspect ratio at short times. The dashed lines show the fits of an exponential decay function to the data. **f**, The viscosity-to-surface-tension ratio, *η/γ*, obtained from droplet coalescence measurements in **e** for the PrLD (*N* = 18 droplets) and PrLD-SAMWT (*N* = 33) condensates.

By contrast, the fluorescence intensity *I*(*t*) of bleached SAMWT fibers remains unrecoverable for over 10 minutes, suggesting that the SAMWT condensate behaves like a solid with little diffusive molecules. The remarkably different material properties between the two condensates can be traced back to their distinct molecular interaction. The interactions in PrLD condensate are weak and relatively isotropic due to its intrinsically disordered structure [30, 31]. SAMWT, on the other hand, is a self-oligomerizing domain, which prefers to form polymeric filaments. These polymeric filaments can further bundle together to form visible fibers or crystals [34, 35].

### Gel-like mechanical behavior of PrLD-SAMWT condensates

The PrLD-SAMWT condensate stays at an intermediate state between liquid-like PrLD condensate and solid-like SAMWT condensate. It retains a droplet shape (Fig. 1**a**, bottom panel), but its FRAP curve reveals a significantly hindered molecular diffusion compared to PrLD (Figs. 1**b** and 1**c**) [21]. Furthermore, the FRAP curve *I*(*t*) for PrLD-SAMWT in Fig. 1**c** does not agree well to the diffusion-driven model that works for PrLD [21] (see SI Section I.D).

We then conducted droplet coalescence measurements to examine its fluid property. As shown in Fig. 1**d**, PrLD droplets fuse into a single spherical droplet within seconds when brought into contact with each other, whereas PrLD-SAMWT droplets take more than an hour to merge. By fitting the fusion curve to an exponential decay function (Fig. 1**e**), we find the characteristic fusion time *τ*_dc_ = 0.22 s for PrLD and *τ*_dc_ = 2168 s for PrLD-SAMWT (see SI Section I.G). This fusion time is determined by [45–49], *τ*_dc_ ≃ *R*_*d*_*η/γ*, where *γ* is the surface tension, *η* is the viscosity, and *R*_*d*_ is the radius of the droplet. Using this equation, we find *η/γ* = 0.034 ± 0.015 s/*µ*m for PrLD and *η/γ* = 345± 133 s/*µ*m for PrLD-SAMWT (Fig. 1**f**). The addition of SAM domains renders its flow property nearly 10^4^ times slower than that for PrLD alone.

To further understand the mechanical property of PrLD-SAMWT condensate, we measured its elastic modulus *E* using an atomic force microscope (AFM) supplemented with a colloidal probe (radius *R*_*p*_ = 5-10 *µ*m) (Fig. 2**a**) or a plane probe (see Fig. S4 and SI Section I.F). We conducted AFM force indentation measurements by compressing a PrLD-SAMWT droplet (radius *R*_*d*_ = 5-10 *µ*m) with the probe at a fixed loading speed *v* = 5 *µ*m/s (Fig. 2**a**(i)), from which we obtained the force indentation curve *F* (*δ*). As shown in Fig. 2**b**, the PrLD-SAMWT condensate is much stiffer than PrLD, as it requires a much larger force to achieve the same indentation.

**FIG. 2.**
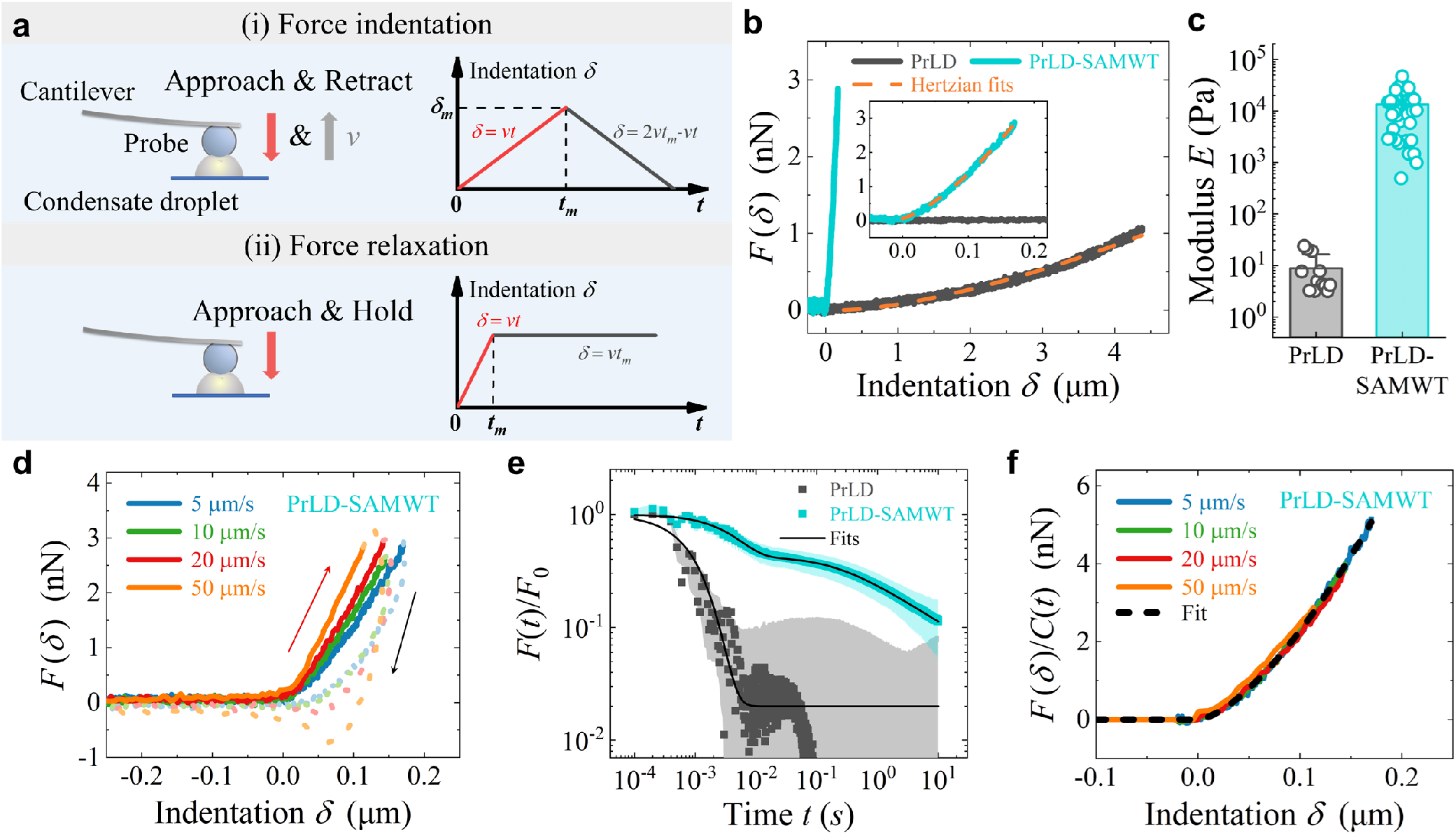
Mechanical characterization of PrLD and PrLD-SAMWT condensates. **a**, Illustration of AFM operation for (i) force indentation measurements, in which the probe approaches and compresses the droplet at a given loading speed *v*, and then retracts from the droplet at the same speed without halting, and for (ii) force relaxation measurements, in which the probe compresses the droplet at a high loading speed *v*, and then holds at a constant indentation *δ* = *vt*_*m*_ on the droplet. The right panels show the resulting indentation *δ*(*t*) as a function of time *t*. **b**, Comparison of the measured force indentation *F* (*δ*) between PrLD and PrLD-SAMWT condensates in advancing direction at a loading speed *v* = 5 *µ*m/s. Inset shows a magnified view of *F* (*δ*) at small indentations. The dashed lines show the Hertzian fits to the data. **c**, Apparent modulus *E* obtained for PrLD (*N* = 12 droplets) and PrLD-SAMWT (*N* = 53) condensates. **d**, Measured *F* (*δ*) in the advancing (colored solid lines) and receding (colored dashed lines) directions made at different loading speeds *v* on one PrLD-SAMWT droplet. **e**, Measured force relaxation, *F* (*t*)*/F*_0_, normalized by the initial force *F*_0_ at time *t* = 0 for PrLD (black squares, *N* = 12) and PrLD-SAMWT (cyan squares, *N* = 50) condensates made at the loading speed *v* = 100 *µ*m/s. The shaded areas indicate the standard deviations of the force measurements. The solid lines show the fits of Eq. (1) to the data. **f**, Normalized force-indentation curves, *F* (*δ*)*/C*(*t*), in the advancing direction at different loading speeds *v*. The data are taken from **d** with the same color codes. The black dashed line shows a fit of Eq. (3) to the data with a fitting parameter *E*_0_ = 24, 012 Pa.

By fitting the measured *F* (*δ*) to the Hertz contact model [18, 50], *F* (*δ*) = (16*/*9)*ER*^1*/*2^*δ*^3*/*2^, where *R* = 1*/*(1*/R*_*p*_ + 1*/R*_*d*_) is the reduced radius, we obtained the apparent elastic modulus *E* = 13, 428± 9, 642 Pa for PrLD-SAMWT (Figs. 2**b** and 2**c**), which is over 1,500 times larger than that for PrLD (*E* = 8.8 ±7.7 Pa, Fig. 2**c**). The measurements of *E* and *η/γ* collectively demonstrate the distinct material properties of PrLD-SAMWT condensate.

### Relaxation modulus *E*(*t*) reveals emergence of a percolated network in PrLD-SAMWT condensate

Figure 2**d** shows the force-indentation curve *F* (*δ*) for a single PrLD-SAMWT droplet at four different loading speeds *v*. The measured approaching curves of *F* (*δ*) (colored solid lines) appear to be of Hertzian type and reveal a strong dependence on *v*. The droplet are stiffer at a higher speed as it requires a larger force to achieve the same indentation. As will be shown below, this speed-dependent *F* (*δ*) is a hallmark of viscoelasticity for protein condensates that exhibit a mixture of fluid- and solid-like behaviors across multiple spatial and temporal scales [18, 51, 52]. To characterize the viscoelastic behavior of PrLD-SAMWT condensate, we conducted additional force relaxation measurement (Fig. 2**a**(ii)). Under this AFM operation, an almost instantaneous (∼ 1 ms) and constant indentation *δ* is imposed onto the droplet, and then the AFM records the resulting force decay *F* (*t*) over a 5-decade time span from 0.1 ms to 10 s during the hold stage [18].

Figure 2**e** shows the normalized force *F* (*t*)*/F*_0_ with *F*_0_ being the initial force at time *t* = 0 over *N* PrLD-SAMWT and PrLD droplets. The force acting on PrLD relaxes rapidly to a baseline near zero, whereas the measured *F* (*t*) for PrLD-SAMWT displays a much longer and slower decay. Both sets of data can be well described by a multiscale relaxation model [18, 53]:

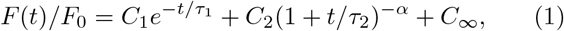

where *C*_1_, *C*_2_ and *C*_*∞*_ are the weighting factors of the three relaxation modes, *τ*_1_, *τ*_2_ are the corresponding relaxation times of the two decay modes, and *α* is the power-law exponent. The solid lines in Fig. 2**e** show the best fits of Eq. (1) to the data. The force relaxation of PrLD-SAMWT condensate has three modes: an initial exponential decay at short times *t* with *C*_1_ = 0.61 ± 0.07 and *τ*_1_ = 4.7 ± 1.5 ms, and a long-time power-law decay with *C*_2_ = 0.34 ± 0.08, *τ*_2_ = 0.48 ± 0.40 s and *α* = 0.59 ± 0.24, and the residue term *C*_*∞*_ = 0.05 ± 0.06, which does not change with time *t*. By contrast, the force relaxation of PrLD condensate contains only an exponential decay with *C*_1_ = 0.98 ± 0.10 and *τ*_1_ = 0.90 ± 0.29 ms, and a small value of *C*_*∞*_ = 0.02 ± 0.10. The absence of long-time power law relaxation explains why the measured *F* (*t*) decays so quickly to a near-zero baseline.

Recent AFM studies of force relaxation of a protein condensate [18] and living cells [53] revealed that the exponential relaxation at short times is caused by the diffusive relaxation of mobile proteins. The power-law decay at long times is caused by a slow relaxation of deformed immobile protein network in the condensate. Figure 2**e** thus indicates that a protein network is formed in the PrLD-SAMWT condensate, which sustains external stresses and relaxes by structural rearrangements of the network. The residual stress *C*_*∞*_ represents a persistent elastic response from the network.

For a fixed indentation *δ*, the force relaxation *F* (*t*) is proportional to the time-dependent Young’s modulus *E*(*t*) of the condensates, which takes the form [18, 53]:

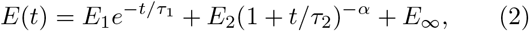

where *E*_1_, *E*_2_ and *E*_*∞*_ are the modulus components of three relaxation modes, defined as *E*_*i*_ = *C*_*i*_*E*_0_ (*i* = 1, 2, *∞*). Here *E*_0_ is the initial modulus at *t* = 0 with *E*_0_ = *E*_1_ + *E*_2_ + *E*_*∞*_.

With the relaxation modulus *E*(*t*), the force indentation relation in the original Hertz contact model is modified to contain a time-dependent factor *C*(*t*) and is given as [18, 52–54],

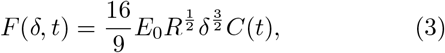

where *t* = *δ/v* and *C*(*t*) = *C*(*t*; *C*_1_, *C*_2_, *C*_*∞*_, *τ*_1_, *τ*_2_, *α*) is a time-dependent function that can be determined from the measured *F* (*t*)*/F*_0_ in Eq. (1) [18]. As shown in Fig. 2**f**, the normalized plot of *F* (*δ*)*/C*(*t*) for the four approaching curves obtained at different loading speeds *v* collapse into a single master curve, which is well described by Eq. (3) with a single fitting parameter *E*_0_ = 24, 012 Pa for PrLD-SAMWT (see Table S1 for all fitting parameters). Our results thus demonstrate that the observed speed-dependence in *F* (*δ*) is caused by the relaxation of *E*(*t*) and that our multi-scale modeling of *E*(*t*) in Eq. (2) captures the essential physics.

### Complex shear modulus *G*^*\**^(*f*) reveals power-law rheology of PrLD-SAMWT condensate

The complex shear modulus, *G*^*\**^(*f*) = *G*^*′*^(*f*) + *iG*^*′′*^(*f*), is an alternative way to characterize the viscoelastic properties of materials in the frequency domain [51, 52]. The real part, the storage modulus *G*^*′*^(*f*), is used to describe material’s elastic response and the ability to withstand elastic deformation. The imaginary part, the loss modulus *G*^*′′*^(*f*), quantifies the energy dissipation during deformation. Figure 3**a** illustrates the AFM setup used to measure *G*^*\**^(*f*). By imposing a sinusoidal indentation on a condensate droplet, the AFM captures the amplitude and phase shift of the force response, from which the shear modulus *G*^*\**^(*f*) is determined [52] (see Fig. S6 and SI Section II.A).

**FIG. 3.**
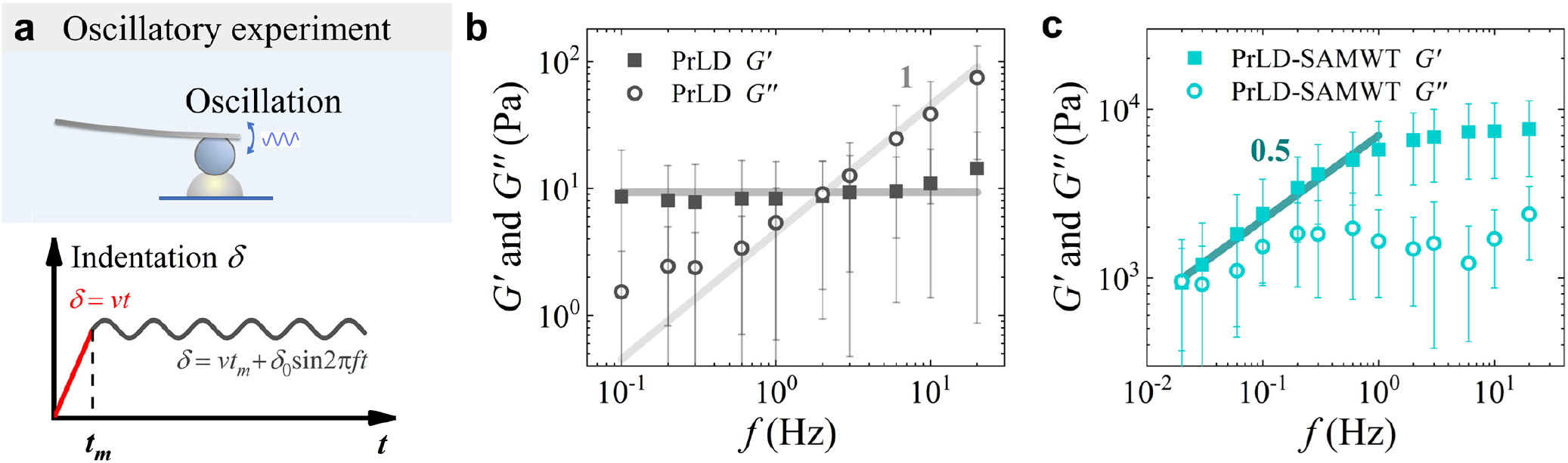
Complex shear modulus *G*^*\**^(*f*) of PrLD and PrLD-SAMWT condensates. **a**, Schematic illustration of AFM setup for shear modulus measurement. The AFM probe imposes a sinusoidal indentation, *δ*(*t*) = *vt*_*m*_ + *δ*_0_ sin(2*πft*), on a condensate droplet after loading. Here *vt*_*m*_is the mean value of indentation, *δ*_0_ is the amplitude of oscillation, and *f* is the oscillation frequency. **b**, Measured storage modulus *G*^*′*^(*f*) and loss modulus *G*^*′′*^(*f*) for PrLD condensate (*N* ≥23 droplets). The error bars show the standard deviation of measurements. The gray lines show the fits of Eq. (4) to the data. **c**, Measured *G*^*′*^(*f*) and *G*^*′′*^(*f*) for PrLD-SAMWT condensate (*N ≥* 19 droplets). The dark cyan line indicates the power-law, *G*^*′*^(*f*) ∝ *f* ^0.5^.

Figure 3**b** shows the obtained moduli *G*^*′*^ and *G*^*′′*^ for PrLD condensate. The measured *G*^*′*^ is almost constant independent of frequency *f* and *G*^*′′*^ is approximately a linear function of *f* . This behavior fits well to the Kelvin-Voigt (KV) model [51, 52], which consists of a spring element with modulus *G*_0_ and a damping element with viscosity *η* connected in parallel. The KV model gives rise to a complex modulus:

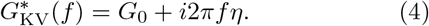

By fitting Eq. (4) to the data, we obtain *G*_0_ = 9.3 Pa and *η* = 0.72 Pa·s for PrLD condensate.

As shown in Fig. 2**e**, the relaxation modulus *E*(*t*) for PrLD condensate is well described by a single exponential decay, which is a hallmark of Maxwell-like fluids [18, 51, 52]. This conclusion is based on the assumption that the viscoelastic response of a condensate droplet to AFM indentation comes predominantly from its bulk viscoelasticity, while contributions from surface effects are negligible. When the surface tension cannot be disregarded, AFM indentation simultaneously deforms both the bulk interior and the interfacial area of the droplet. As a result, the droplet surface tension gives rise to a small value of *C*_*∞*_ in Fig. 2**e** and a constant value of *G*^*′*^ in Fig. 3**b** [48, 55, 56] (see Fig. S7 and SI Section II.B). This result explains why the PrLD condensate fits well with the KV model and behaves as a Maxwell fluid. With the measured viscosity *η* and viscosity-to-surface-tension ratio *η/γ* in Fig. 1**f**, we calculate the surface tension, *γ* = 21 *µ*N/m, for PrLD droplets.

As shown in Fig. 3**c**, PrLD-SAMWT condensate has a drastically different shear modulus *G*^*\**^(*f*). First, its overall magnitude is several hundred times larger than that for PrLD. Second, it shows a very different frequency-dependence. At low frequencies, both *G*^*′*^(*f*) and *G*^*′′*^(*f*) go as a power law, *G*^*′*^(*f*)∝ *f* ^*α*^ and *G*^*′′*^(*f*) ∝ *f* ^*α*^ with *α* ≃ 0.5. When *f* becomes larger than a crossover frequency *f*_*c*_ ≃2 Hz, *G*^*′*^(*f*) saturates at a plateau value *G*_*c*_ and *G*^*′′*^(*f*) witnesses a slight decrease over *f* . The appearance of power-law rheology for *G*^*\**^ (*f*) at low frequencies (or for *E*(*t*) at long times) signify the emergence of a percolated network in PrLD-SAMWT condensate, which is absent in PrLD condensate. Similar power-law rheology was also observed in a networked multivalent protein condensate [18].

At high frequencies (> *f*_*c*_), the percolated network behaves like an elastic spring with a dominant modulus *G*_*c*_, which resists imposed oscillations. At lower frequencies (< *f*_*c*_), both *G*^*′*^(*f*) and *G*^*′′*^(*f*) decrease as the network structure becomes dynamically weaker due to binding/unbinding activities of bonds of the network. This transient nature of cross-links in protein networks gives rise to an unusual force relaxation that allows the network to reorganize or flow at long times. The model of cross-link-governed dynamics by Broedersz *et al*. [57] provided a mechanical description to this dynamic behavior. It predicted a power-law relaxation with *α* = 1*/*2, in agreement with our experimental observations. Furthermore, the power-law relaxation starts at time *t* ≳ *τ*_off_, where *τ*_off_ is a characteristic time for a cross-linker to unbind. As shown in Fig. 3**c**, the crossover frequency *f*_*c*_ ≃ 2 Hz gives a characteristic time, *τ*_off_ ≃ 1*/f*_*c*_ ≃ 0.5 s, which agrees with the power-law relaxation time, *τ*_2_ = 0.48 s, obtained in Fig. 2**e**.

### Altered SAM-SAM interactions by mutation affect SAM oligomerization and protein mobility in PrLD-SAM condensates

The drastic mechanical differences between PrLD and PrLD-SAMWT condensates demonstrate the critical role of SAM in forming a percolated network inside the condensate. To further understand the transitional behavior of network formation, we systematically vary the SAM-SAM interactions by introducing different single amino-acid mutations to the SAM domain. By replacing the wild type methionine (M), which is located at the 56th amino acid in the SAM domain, by other amino acids, including glutamic acid (E), alanine (A), arginine (R), cysteine (C), histidine (H), leucine (L) and lysine (K), we generate a series of PrLD-SAM proteins carrying specific mutations (see SI Section I.A for sample preparations). Hereafter, we label these 7 mutant proteins as ME, MA, MR, MC, MH, ML and MK. The PrLD-SAMWT is labeled as WT.

To characterize the polymerization capability of SAM mutants, we used size exclusion chromatography coupled with multi-angle light scattering (SEC-MALS assay) to measure the degree of oligomerization of PrLD-SAM (Fig. 4**b**). Since the condensate volume is very limited, we used uniform PrLD-SAM mutant solutions with solubility tags on at a given concentration (100 *µ*M) for SEC-MALS. When a PrLD-SAM solution flows through the porous medium column, protein oligomers are separated based on their hydrodynamic size. The eluted solution is subsequently analyzed by light scattering to measure the molecular weight *M*_w_ of the oligomers and apparent oligomerization number *N*_oli_ based on the measured differential refractive index (dRI) (Figs. 4**c** and 4**d**).

**FIG. 4.**
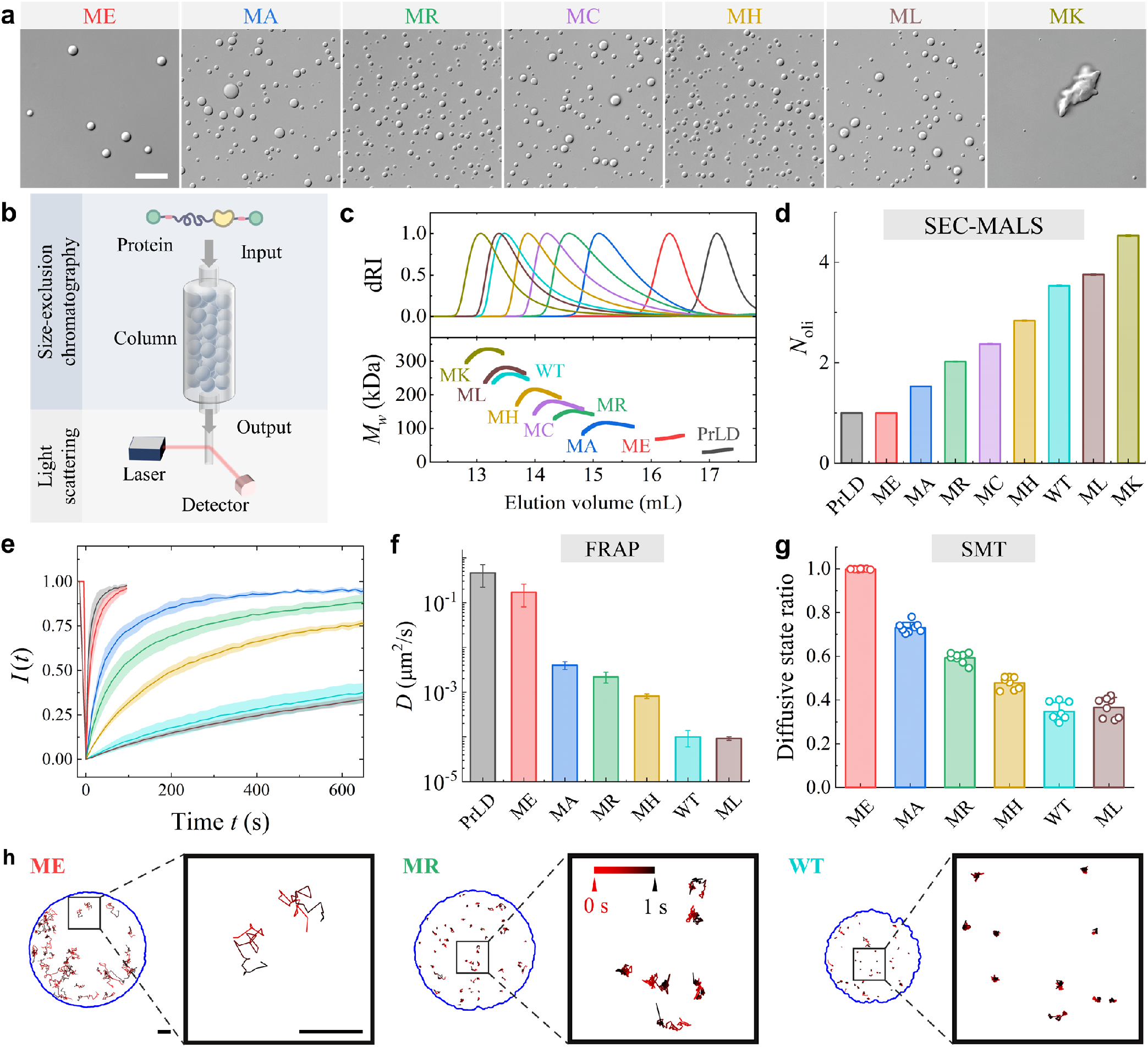
Molecular characterization of PrLD-SAM condensates. **a**, Morphology of phase-separated condensate of various PrLD-SAM mutants. Scale bar: 20 *µ*m. **b**, Illustration of experimental setup for size exclusion chromatography coupled with multi-angle light scattering (SEC-MALS assay). **c**, Measured differential refractive index (dRI, upper panel) and molecular weight *M*_w_ (lower panel) of eluted oligomers in PrLD and PrLD-SAM solutions with solubility tags. **d**, Obtained oligomerization number, *N*_oli_, of eluted oligomers in PrLD and PrLD-SAM solutions. **e**, Measured FRAP curves for PrLD, ME, MA, MR, MH, WT and ML condensates (*N* = 6, 7, 6, 14, 7, 9, and 20 droplets, respectively). The color code used is the same as that in **d**. The solid lines show the mean values and the shaded areas indicate the standard deviations of measurements. **f**, Diffusion coefficient *D* obtained from the FRAP measurements in **e** for PrLD and PrLD-SAM condensates. The error bars show fitting uncertainties. **g**, Diffusive state ratio *σ* for WT and 5 mutants from ME to ML, measured by single molecule tracking (SMT) (*N* = 7, 8, 8, 8, 8, and 8, respectively). **h**, Representative single-molecule trajectories of fluorescent-labeled proteins in ME, MR, and WT condensates over a period of 1 second. Magnified views of protein trajectories are shown on the right of each image pair. Scale bars: 1 *µ*m. The protein trajectories start in red (0 s) and end in black (1 s). Single-molecule trajectories in MA, MH and ML condensates are shown in Fig. S3. The FRAP data for PrLD and WT in **e** are adopted from Fig. 1**c** as a control for comparing the results with mutants.

As shown in Fig. 4**d**, the PrLD and ME are monomeric with their apparent oligomerization numbers *N*_oli_ = 1. Starting from MA, the mutant’s oligomerization number *N*_oli_ increases gradually from 1 to 4.5, suggesting an increased SAM-SAM binding strength and polymerization capability from MA to MK. The actual values of *N*_oli_ for PrLD-SAM mutants in the condensate are expected to increase even more, because the solubility tags may limit SAM-SAM polymerization by their effect to enhance the affinity of PrLD-SAM mutants with water. Nonetheless, the SEC-MALS assay provides a useful measure of relative polymerization capability of PrLD-SAM, as the effect of solubility tags on polymerization remains the same for all mutants. The seven mutants thus produce a smooth gradient of interaction strength from weak to strong (out of 20 available amino acids, see Fig. S1 and SI Section I.C), covering the transition range of material properties from liquid-like condensates (e.g., PrLD) to gel-like condensates (e.g., PrLD-SAMWT).

Upon tag cleavage, most PrLD-SAM mutants phase-separate into spherical droplets, while MK appears as large aggregates with irregular shapes, as shown in Fig. 4**a**. This is plausibly because the SAM-SAM interactions in MK is so strong that SAMMK domains polymerize and form long fibers, which further bundle together and prevent MK from forming spherical droplets, similar to that for SAMWT aggregates (Fig. 1**a**, bottom panel). Due to its irregular shape, we did not conduct further molecular and mechanical measurements for MK.

We next examine molecular diffusion of PrLD-SAM mutants in the condensates by FRAP. As shown in Fig. 4**e** (and Fig. S2), the recovery of fluorescence intensity *I*(*t*) decreases with increasing SAM-SAM interactions. By fitting the FRAP curve to a diffusion-driven model, we find that the fitted values of averaged protein diffusion coefficient *D* also decrease with increasing SAM–SAM interactions (Fig. 4**f**). MC is not included in the FRAP assay because the cysteine-conjugated fluorescence labeling was found to disturb the SAM-SAM interactions of MC (See SI Section I.B).

Single molecule tracking (SMT) was also used to study the diffusion dynamics of fluorescent-labeled individual PrLD-SAM molecules [21] (See SI Section I.E). As SAM-SAM interaction increases, protein trajectories over a fixed period (1 s) become more localized, indicating a decreased molecular diffusion (see Fig. 4**h** for three examples and three more cases are shown in Fig. S3). For condensates from MA to ML, the protein motion undergoes a continuous switch between a mobile diffusive state, in which molecules can travel over a long distance, and a transiently immobile state, in which molecules jiggle around in a confined space [21]. The diffusive state ratio *σ*, defined by the fraction of time spent in the diffusive state, is found to decrease gradually with increasing SAM-SAM interactions (Fig. 4**g**). As SAM interaction becomes stronger, longer SAM polymer chains and larger SAM clusters form in the condensate. As a result, the PrLD-SAM molecules become less diffusive with decreasing *D* and *σ*.

### Material properties of PrLD-SAM condensates feature a sharp rigidity percolation transition

In contrast to the gradual changes in molecular properties (such as *N*_oli_, *D* and *σ*), the material properties of PrLD-SAM condensates reveal a sharp transition as SAM-SAM interactions increase. Figure 5**a** shows the obtained apparent modulus *E* for PrLD and PrLD-SAM condensates (see Fig. S9 for force indentation curves). The eight samples can be divided into three groups based on their values of *E*. First, the measured *E* for ME is approximately the same as that for PrLD. Second, the modulus *E* for MA, MR, MC, and MH remain approximately the same but several times larger than that for PrLD. Finally, the modulus *E* for ML is of the same order as that for WT, which is 269 times larger than that for MH. Further-more, the measured viscosity-to-surface-tension ratio *η/γ* for PrLD and PrLD-SAM condensates reveals a similar trend as *E* (Fig. 5**b** and Fig. S5). The obtained *η/γ* for WT is 427 times larger than that for MH. The remarkable changes in condensate stiffness and fluidity suggest that the condensate mechanics undergoes a sharp transition when the SAM-SAM interactions are increased from MH to WT.

**FIG. 5.**
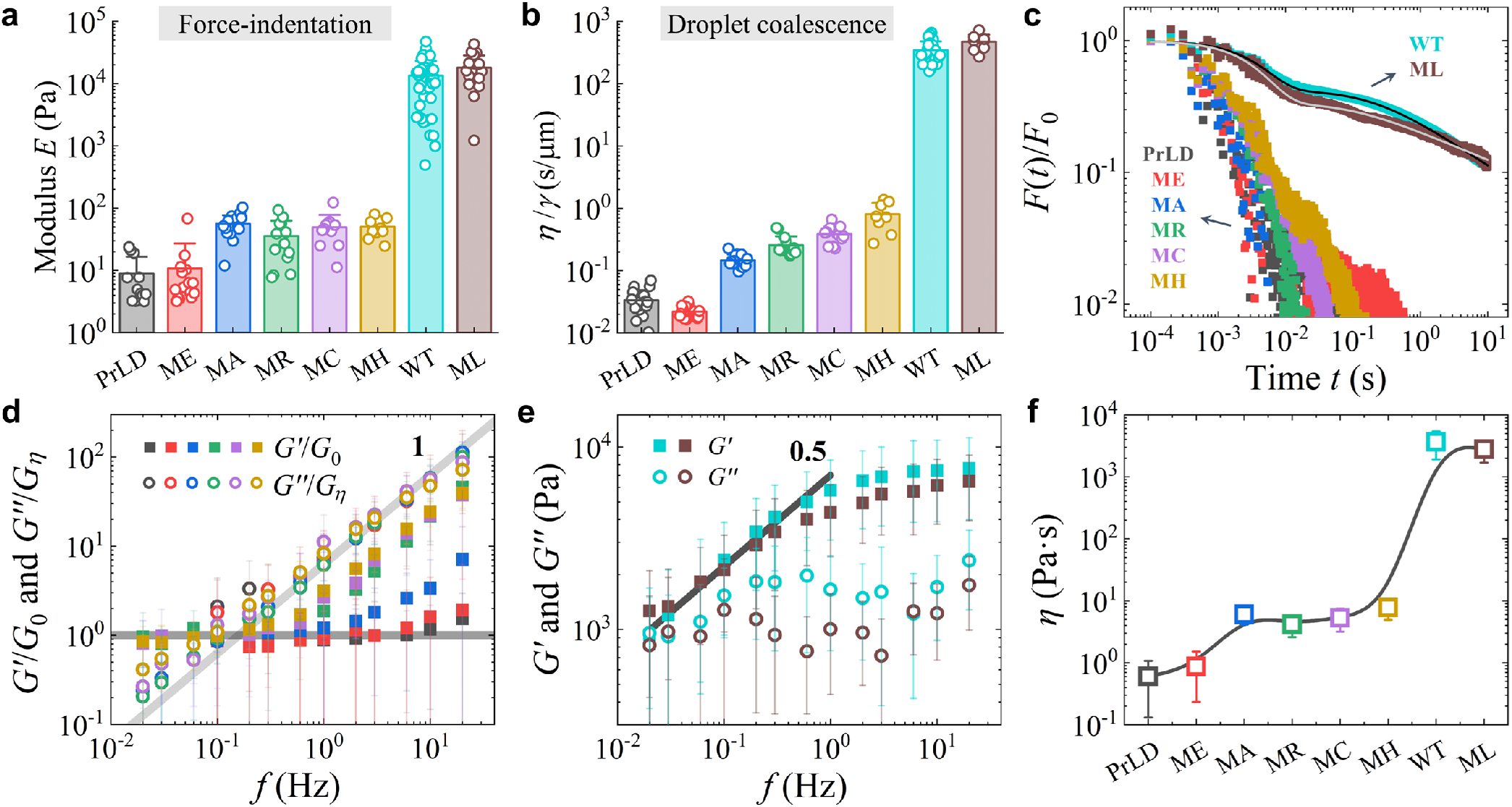
Characterization of a sharp rigidity percolation transition in PrLD-SAM condensates. **a**, Apparent modulus *E* for PrLD and PrLD-SAM condensates obtained from the force indentation measurements (*N* =12, 15, 17, 13, 11, 10, 53, and 18 droplets for PrLD, ME, MA, MR, MC, MH, WT, and ML, respectively). **b**, Viscosity-to-surface-tension ratio *η/γ* obtained from droplet coalescence measurements for PrLD and PrLD-SAM condensates (*N* = 18, 22, 14, 15, 12, 8, 33, and 8, respectively). **c**, Normalized force relaxation curves, *F* (*t*)*/F*_0_, obtained for PrLD and PrLD-SAM condensates (averaged over *N* = 12, 13, 17, 13, 11, 15, 50, and 16 droplets, respectively). The two upper solid lines show the fits of Eq. (1) to the WT and ML data. **d**, Normalized storage modulus *G*^*′*^(*f*)*/G*_0_(closed squares) and loss modulus *G*^*′′*^(*f*)*/G*_*η*_ (open circles) as a function of frequency *f* for PrLD, ME, MA, MR, MC and MH condensates (*N*≥ 23, 21, 17, 10, 11 and 14, respectively). The color code used is the same as that in **c**. The two gray solid lines indicate the power laws, *G*^*′*^(*f*)*/G*_0_ ∼*f* ^0^ and *G*^*′′*^(*f*)*/G*_*η*_ ∼2*πf* ^1^, respectively. The error bars show the standard deviations of measurements. **e**, Measured storage modulus *G*^*′*^(*f*) (closed squares) and loss modulus *G*^*′′*^(*f*) (open circles) for WT (cyan, *N* ≥ 19) and ML (brown, *N* ≥ 17) condensates. The black solid line indicates the power law *G*^*′′*^(*f*)∼ *f* ^0.5^. The error bars show the standard deviation of measurements. **f**, Obtained viscosity *η* (open squares) for PrLD and PrLD-SAM condensates. The error bars show the standard deviation of measurements. The solid line is a B-spline fit for visual guidance. The data for PrLD and WT in **a**-**e** are adopted from Figs. 1**f**, 2**c**, 2**e**, 3**b**, and 3**c** as a control for comparison.

To further investigate the mechanical changes in PrLD-SAM condensates, we conducted force relaxation and oscillatory measurements. Figure 5**c** shows the normalized force relaxation curves *F* (*t*)*/F*_0_ for PrLD and PrLD-SAM condensates. The eight force relaxation curves can be divided into two groups. The first group, including PrLD, ME, MA, MR, MC and MH, exhibits a rapid and exponential-like force relaxation, suggesting that their relaxation behavior resembles Maxwell-like fluids. The second group consisting of WT and ML features a much slower, power-law relaxation at long times, which is a hallmark of network relaxation (see Table S1 for all fitting parameters). This sharp transition, which occurs between MH and WT, is associated with the emergence of a percolated protein cluster/network when the cross-link density (or protein connectivity) exceeds the percolation threshold [23, 26].

We also measured the complex shear modulus *G*^*\**^(*f*) for PrLD, ME, MA, MR, MC and MH condensates (Fig. 5**d**). The measured *G*^*\**^(*f*) at low frequencies for the six condensates can all be well described by the KV model, as shown in Eq. (4), with different values of *G*_0_ and *η*. In Fig. 5**d**, we normalize the measured *G*^*′*^(*f*) by *G*_0_ and *G*^*′′*^(*f*) by *G*_*η*_ = *ηf*_0_ with *f*_0_ = 1 Hz being a fixed reference frequency, so that all the storage and loss modulus curves overlap on two master curves, *G*^*′*^(*f*)*/G*_0_ ∼ *f* ^0^ and *G*^*′′*^(*f*)*/G*_*η*_ ∼ 2*πf* ^1^, at low frequencies (*f* ≲ 0.6 Hz). At higher frequencies, the normalized storage modulus *G*^*′*^(*f*)*/G*_0_ for MA, MR, MC and MH starts to increase with *f* . This enhanced elastic response is closely linked to the enhanced SAM interactions and increased oligomerization of PrLD-SAM chains (see Fig. 4**d**). With increasing *N*_oli_, the condensates change from Newtonian-like fluids, typical for small monomeric molecules, to polymeric solutions, which manifests as Maxwell-like stiffening in the measured *G*^*′*^(*f*). Similar changes were also found for PGL-3 condensate [48, 58].

The measured *G*^*\**^(*f*) for ML is similar to WT, showing an elastic-dominant behavior at high frequencies and distinct power-law rheology, *G*^*′*^(*f*) ∼ *f* ^0.5^, at low frequencies (Fig. 5**e**). The shared features between ML and WT imply that a percolated network is formed inside both condensates and the condensate mechanics may saturate after the percolation transition. The mechanical characterization of PrLD-SAM condensates clearly reveals a rigidity percolation transition when the SAM-SAM interactions are increased across a particular threshold between MH and WT. Beyond the threshold, a percolated network structure is formed inside the condensates, which results in a drastic stiffening of the condensate mechanics. This sharp rigidity transition is not parallel to the molecular properties of the condensates, revealing that network formation is indeed emergent at the condensate level.

From the measured viscosity-to-surface-tension ratio *η/γ* in Fig. 5**b** and loss modulus *G*^*′′*^(*f*) in Figs. 5**d** and 5**e**, we calculate the absolute value of the condensate viscosity *η* and surface tension *γ* separately, as shown in Fig. 5**f** and Fig. S8. For WT and ML, we used an empirical relation [18, 58], *η* ≃ *G*_*c*_*τ*_2_, to estimate their viscosity. The obtained condensate viscosity *η* displays the same characteristic three-tier transition as the apparent modulus *E* (Fig. 5**a**). By comparison, the droplet surface tension *γ* decreases more gradually with increasing SAM-SAM interactions.

### Critical behavior of the rigidity percolation transition in binary MH-WT condensates

To examine the critical behavior of the rigidity percolation transition occurred between MH and WT condensates, we prepare a series of binary condensates composed of MH and WT proteins with varying molar ratios while keeping the total PrLD-SAM concentration constant. By doping weaker SAM bonds of MH into the percolated network formed by WT, we reduce the effective oligomerization number of SAM chains, introduce heterogeneous SAM-SAM interactions, and thereby fine-tune the network mechanics in the vicinity of the rigidity percolation transition.

Figure 6**a** shows the viscosity-to-surface tension ratio, *η/γ*, obtained for MH-WT condensates with varying WT molar fractions *ϕ*. The measured *η/γ* shows a sharp rise starting at the critical molar fraction *ϕ*_*c*_ ≃ (64±1.5)%, indicating the onset of a distinct transition in the material properties of the binary condensates.

**FIG. 6.**
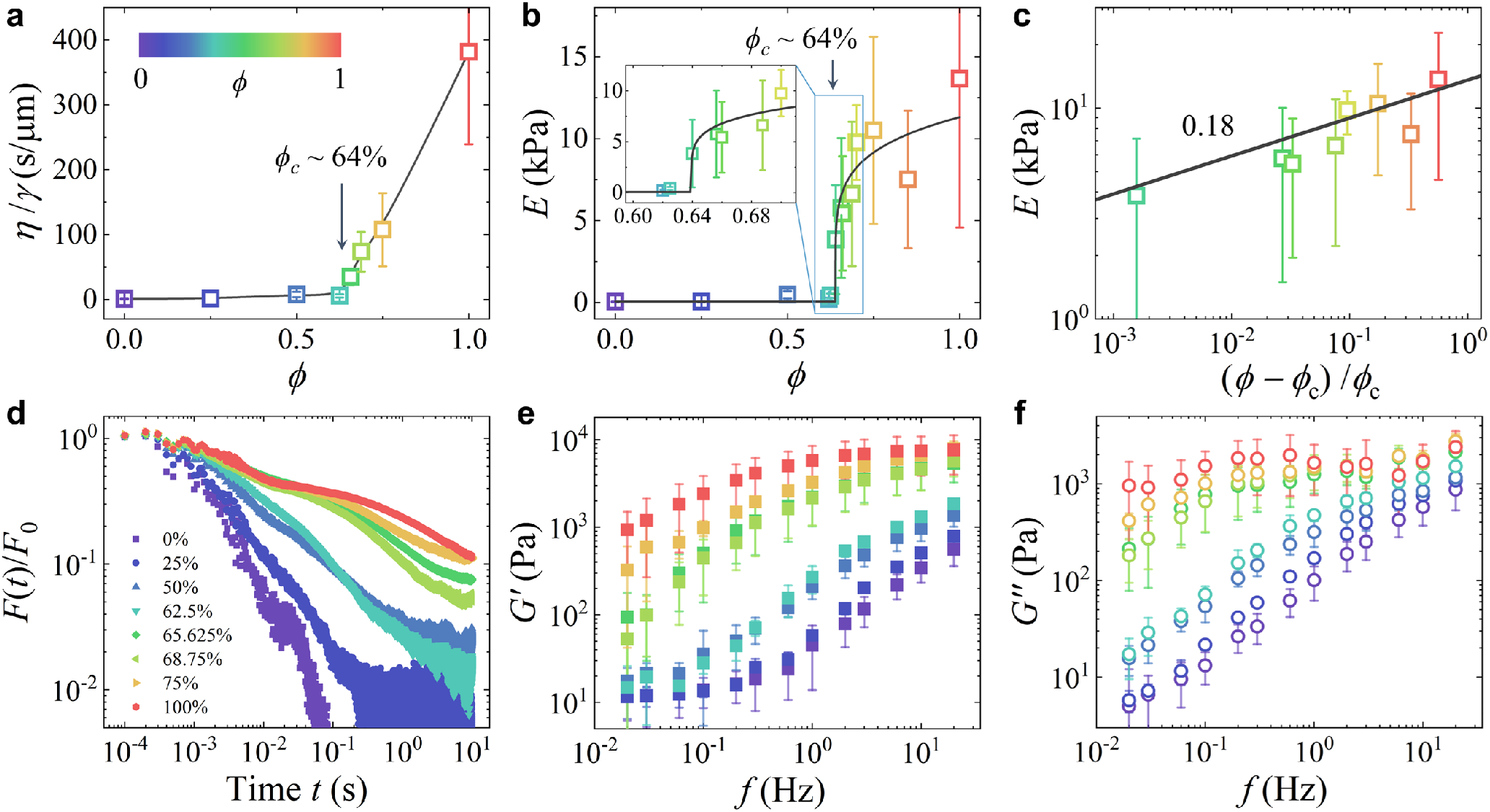
Evidence for criticality in the rigidity percolation of binary MH-WT condensates. **a**, Measured viscosity-to-surface tension ratio *η/γ* as a function of WT molar fraction *ϕ* for the binary condensates. The arrow points to the onset of transition, at which *η/γ* starts to increase rapidly. The black solid line is a spline fit for visual guidance. **b**, Measured apparent modulus *E* as a function of WT molar fraction *ϕ* for the binary condensates. The arrow points to the boxed transition region (enlarged in the inset). The black solid lines indicate the power-law scaling, *E* ∝ (*ϕ −ϕ*_*c*_)^*β*^. **c**, Replot of the measured *E* in the transition region as a function of reduced WT molar fraction, (*ϕ− ϕ*_*c*_)*/ϕ*_*c*_. The black solid line shows a power law fit to the data with the exponent *β* = 0.18± 0.11. **d**, Normalized force relaxation curves, *F* (*t*)*/F*_0_, obtained with different WT molar fractions *ϕ* for the binary condensates. **e** and **f**, Measured storage modulus *G*^*′*^(*f*) (**e**) and loss modulus *G*^*′′*^(*f*) (**f**) for the binary condensates with different WT molar fractions *ϕ*. The color codes defined in **a** and **d** are the same and apply to all the panels in **a**–**f**. The error bars show the standard deviations of measurements. The data for MH (*ϕ* = 0) and WT (*ϕ* = 1) in **a**–**f** are adopted from Figs. 5**a**–**e** as a control for comparison.

A similar transition is observed in the apparent modulus *E* of the binary condensates, as shown in Fig. 6**b**. It is found that the modulus *E* of the binary condensates is well described by a power-law scaling relation [22, 23]

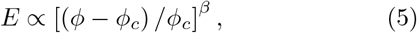

where *β* is the critical exponent. Fitting Eq. (5) to the data in Figs. 6**b** and 6**c** yields a critical molar fraction of *ϕ*_*c*_ = (63.9± 0.6)% and a scaling exponent of *β* = 0.18±0.11. Figures 6**a**–6**c** thus provide clear evidence of critical behavior associated with the rigidity percolation transition in binary MH-WT condensates.

Figure 6**d** presents the normalized force relaxation curves, *F* (*t*)*/F*_0_, for MH-WT binary condensates at varying WT molar fractions *ϕ*. As *ϕ* increases, force relaxation in the binary condensates slows progressively. Notably, a power-law-like relaxation emerges when *ϕ* reaches ∼ 50%, which is lower than the critical molar fraction *ϕ*_*c*_ ≃ 64% determined from the measurements of *E* and *η/γ*. This apparent discrepancy in *ϕ*_*c*_ arises from the dynamic nature of the percolated network and the different timescales probed by the respective experiments. Over time, the protein network can relax stresses through structural rearrangements facilitated by the dynamic binding and unbinding of crosslinks [18, 57], which fluidizes the network, enables macroscopic flow and thereby accounts for the relatively small values of *E* and *η/γ* measured over long timescales. This effective reduction in mechanical strength shifts the percolation transition to higher *ϕ*.

The dynamic nature of the protein network is also evident from the fitted network relaxation time *τ*_2_. Compared with pure WT condensate, the network in binary condensates is more dynamic (shorter *τ*_2_) and more liquid-like (larger exponent *α*). At *ϕ* = 50%, the condensate network exhibits a short unbinding time of *τ*_2_ ≃ 19 ms and a large power-law exponent *α* ≃ 0.8 (see Table S2 for all fitting parameters). As *ϕ* increases, *τ*_2_ lengthens, indicating progressive stabilization of the network structure due to the incorporation of stronger SAM bonds. The amplitude of the relaxation modulus *E*(*t*) also increases with *ϕ*, further highlighting the dynamic character of cross-linking at different timescales.

The measured shear modulus *G*^*\**^(*f*) reveals a similar trend as the relaxation modulus *E*(*t*) for MH-WT binary condensates (Figs. 6**e** and 6**f**). As *ϕ* increases, the mechanical property of binary condensates undergoes a transition from liquid-like to gel-like. The obtained *G*^*′*^(*f*) and *G*^*′′*^(*f*) curves separate into two distinct groups with a large gap in magnitude when *ϕ* crosses *ϕ*_*c*_ ≃ 64%. For *ϕ* < *ϕ*_*c*_, the measured *G*^*′*^(*f*) follows a pattern similar to that for pure MH condensate (*ϕ* = 0%), albeit with modestly higher absolute values, and remains approximately constant at low frequencies. For *ϕ* > *ϕ*_*c*_, the low-frequency slope of the measured *G*^*′*^(*f*) gradually decreases from nearly 1 toward 0.5, reflecting characteristic power-law rheology. This behavior is consistent with the force relaxation results presented in Fig. 6**d**. Overall, Figure 6 demonstrates the critical behavior of the rigidity percolation transition, driven by increasing protein interactions, that underlies the changes in material properties from liquid-like to gel-like.

### Disease-associated mutations in SAM impair network percolation in condensates

In addition to the synthetically generated mutations discussed above, we also examine two pathological mutations in the SAM domain of Shank protein that have been identified in human patients with neurodevelopmental disorders, including developmental delay and behavioral abnormalities [38, 41, 42]. The first is the Shank3-K1670* nonsense mutation, which introduces a premature stop codon and truncates the entire SAM domain (also referred to as Shank without SAM) [41, 42]. Removing the SAM domain from Shank corresponds to removing it from PrLD-SAM in our system, leaving only the PrLD domain. As shown above, PrLD alone exhibits Maxwell fluid behavior without forming network structures. Consequently, loss of SAM domain shifts condensate mechanics toward a more dynamic, liquid-like state.

The second is the Shank2-L1800W missense mutation, which replaces leucine (L) at the 1800th residue (the 28th residue of the SAM domain) with tryptophan (W) [38]. To study the effect of this mutation, we generated the PrLD-SAM mutant, PrLD-SAMLW (LW). The SEC-MALS assay shown in Figs. 7**a** and 7**b** reveals that the oligomerization ability of LW mutant is reduced, with a smaller molecular weight and oligomerization number *N*_oli_ compared to WT and MH, indicating a weaker SAM-SAM interaction in the solution. Consequently, the material properties of LW condensate change drastically. As shown in Figs. 7**c** and 7**d**, the measured apparent modulus *E* and viscosity-to-surface-tension ratio *η/γ* for LW are more than two orders of magnitude smaller than those of WT. The FRAP assay for LW also reveals a fast fluorescence recovery to over 90%, as shown in Fig. 7**e**. The liquid-like behavior of LW condensate is expected, as its molecular parameters associated with SAM-SAM interactions fall below the percolation threshold as determined between the MH and WT.

**FIG. 7.**
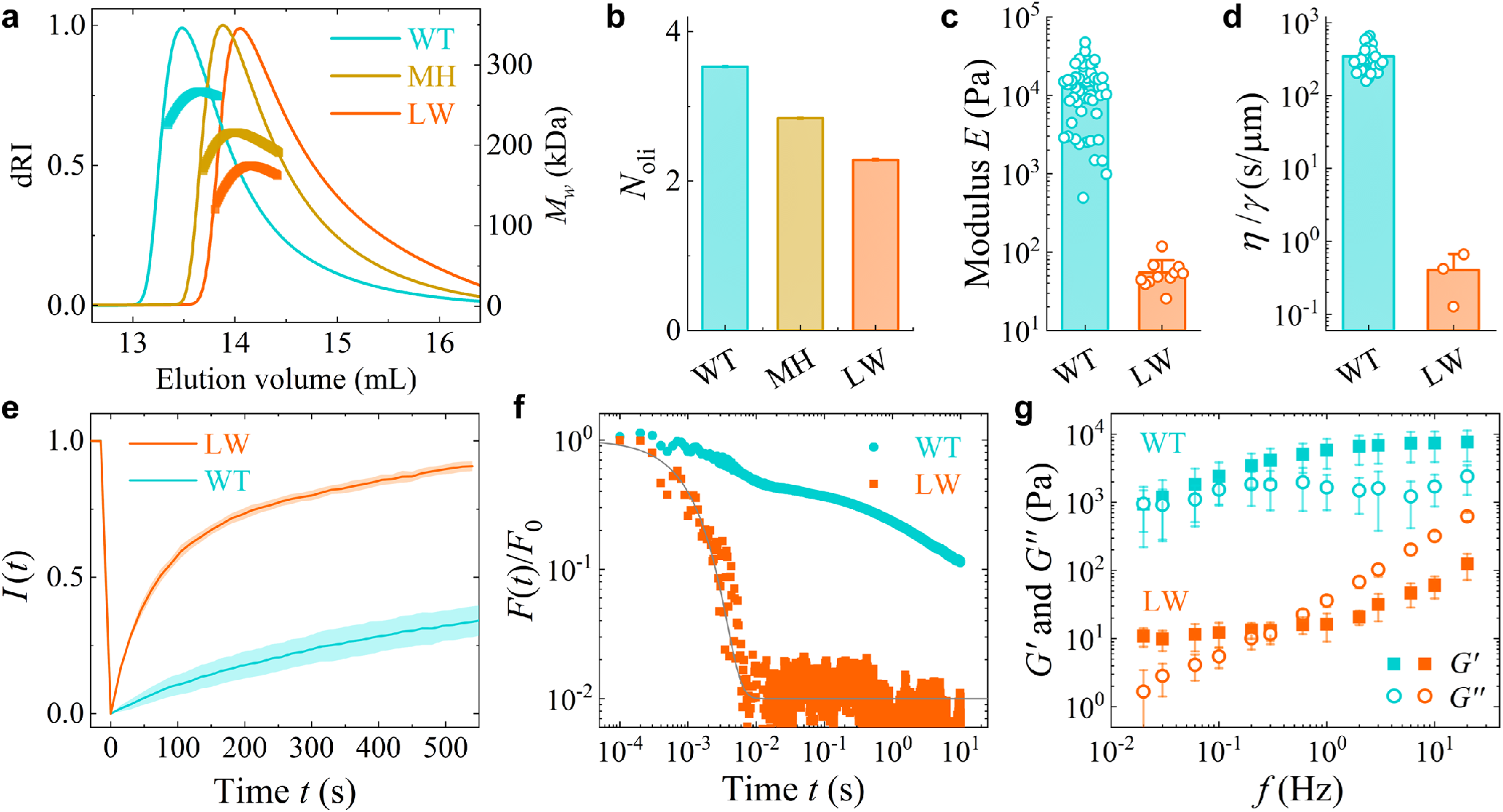
Characterization of disease-associated mutant condensates. **a**, Measured differential refractive index (dRI, thin lines) and molecular weight *M*_*w*_ (thick lines) of WT, MH and LW using the SEC-MALS assay. **b**, Measured apparent oligomerization number *N*_oli_ of WT, MH and LW. **c**, Comparison of apparent modulus *E* between WT (*N* = 53 droplets) and LW (*N* = 11) obtained from force indentation measurements. **d**, Comparison of viscosity-to-surface-tension ratio *η/γ* between WT (*N* = 33) and LW (*N* = 3) obtained from droplet coalescence measurements. **e**, Measured FRAP curves for WT (*N* = 9) and LW (*N* = 6) condensates. The shaded areas indicate the standard deviations of measurements. **f**, Normalized force relaxation curves, *F* (*t*)*/F*_0_, for WT (*N* = 50) and LW (*N* = 11) condensates. The gray solid line shows a fit of Eq. (1) to the data without a power-law decay (i.e. *C*_2_ = 0). **g**, Measured storage modulus *G*^*′*^(*f*) and loss modulus *G*^*′′*^(*f*) for WT (*N≥* 19) and LW (*N≥* 9) condensates. The error bars show the standard deviations of measurements. The data for MH and WT in **a**–**g** are adopted from Figs. 4**c, d, e** and 5**a**–**e** as a control for comparison.

Additional measurements of force relaxation *F* (*t*)*/F*_0_ and complex shear modulus *G*^*\**^(*f*) further confirm that LW condensate behaves like a Maxwell fluid. As shown in Fig. 7**f**, force relaxation in LW condensate is exponentially rapid and decays to a small residual constant *C*_*∞*_ without any detectable power-law tail. This result confirms that LW condensate is well described by the Kelvin-Voigt (KV) model for viscoelasticity without a percolated network. Consistent results are obtained from the measured storage and loss moduli *G*^*′*^(*f*) and *G*^*′′*^(*f*) (Fig. 7**g**). At low frequencies, LW condensate displays a characteristic KV behavior, with a nearly constant *G*^*′*^(*f*) and a linearly increasing *G*^*′′*^(*f*) ∝ *f* . Above 1 Hz, however, *G*^*′*^(*f*) begins to rise markedly, exhibiting Maxwell-like stiffening reminiscent of that seen in MR, MC, and MH condensates. From the above analysis, we find that the two disease-related mutations are all linked to the collapse of percolated network by weakening the SAM-SAM interactions.

It is particularly noteworthy that a single point mutation from leucine to tryptophan in Shank SAM produces dramatic changes in the material properties of the condensate. This effect arises from the mutation-induced shifting across the percolation critical threshold, despite that the mutation causes only minor alterations in the SAM domain’s classical hydrodynamic properties. This observation offers a new perspective on how single-point missense mutations in proteins may drastically alter their interaction networks and thus cause diseases.

## Discussion

The combined AFM-based mesoscale rheology and molecular characterization provide a quantitative molecular and mechanical profiling for PrLD-SAM condensates across a range of protein interaction strengths (see Table. S3 for all parameters). At the molecular level, both the protein oligomerization number *N*_oli_ and immobile protein fraction 1*− σ* increase progressively with stronger SAM-SAM interactions (left panel in Fig. 8**a**). At the condensate level, the material properties of the PrLD-SAM condensates undergo a rigidity transition marked by sharp increases in apparent modulus *E* and viscosity *η* (right panel in Fig. 8**a**). This transition arises from the formation of a percolated protein network, which occurs only when the SAM-SAM interaction strength exceeds a critical threshold between MH and WT.

**FIG. 8.**
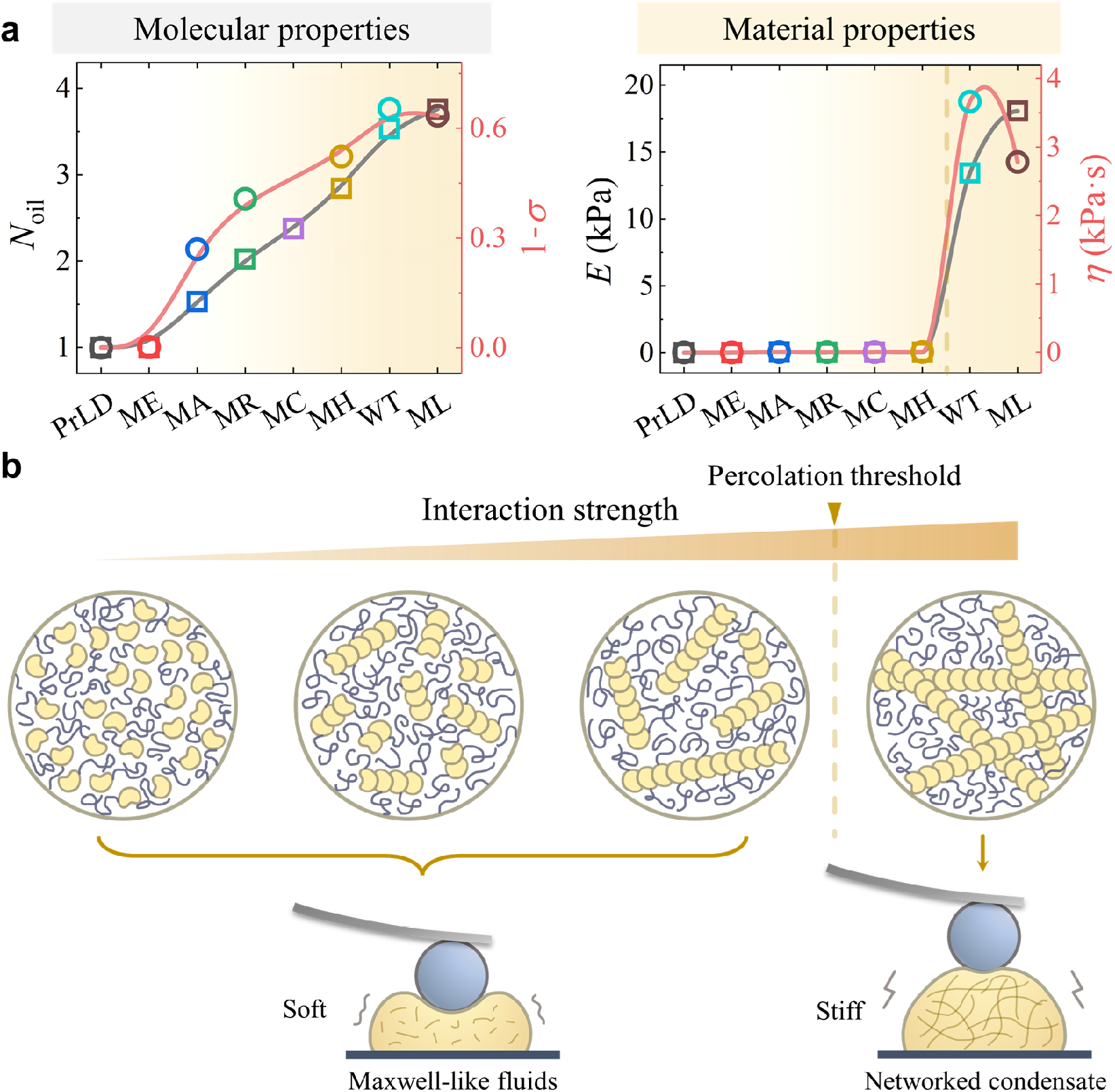
Rigidity percolation transition in PrLD-SAM condensates. **a**, Molecular and mechanical parameters as a function of SAM-SAM interaction strength (increasing from left to right, as indicated by a shade intensity gradient). **Left:** Molecular-level parameters, including the oligomerization number *N*_oli_ (colored squares and grey spline; data from Fig. 4**d**) and the immobile protein fraction 1*− σ* (colored circles and red spline; data from Fig.4**g**), both of which increase gradually with stronger SAM–SAM interactions. **Right:** Material properties, including the apparent modulus *E* (colored squares and grey spline; data from Fig. 5**a**) and viscosity *η* (colored circles and red spline; data from Fig. 5**f**), which exhibit a sharp rigidity percolation transition upon crossing a critical interaction strength (vertical dashed line) between the MH and WT. **b**, Illustration of molecular structural evolution in PrLD-SAM condensates. As the SAM–SAM interaction strength increases, PrLD-SAM chimeras progressively assemble into larger oligomers or polymer chains, leading to greater molecular connectivity within the condensate. Above the percolation threshold, the oligomers or polymer chains spontaneously self-organize into an interconnected protein network spanning the entire droplet. The formation of this percolated network dramatically stiffens the condensate, providing additional mechanical strength against deformation and substantially slowing the mobility of proteins.

Below this threshold, the condensates behave as Maxwell-like (or polymeric) fluids, with their mechanical properties dominated by viscosity and surface tension. The condensates remain soft (fluid-like) because SAM polymerization is weak and cannot form a percolated network, leaving polymeric chains and clusters fluidic and floppy under external deformation (left panel in Fig. 8**b**). Above the threshold, a percolated protein network forms, enabling the condensates to support mechanical loads and exhibit power-law rheology in the relaxation modulus *E*(*t*) and the complex shear modulus *G*^*\**^(*f*) (right panel in Fig. 8**b**). Our results show that network percolation is an emergent collective property of PrLD-SAM and reveal how such a network forms and strengthens with increasing SAM-SAM interactions.

In the vicinity of percolation transition, the condensate rigidity is governed by the power law scaling, *E* ∼ (*ϕ− ϕ*_*c*_)^*β*^, as shown in Eq. (5). Although rigidity percolation has been studied extensively over the past four decades, the predicted values of *β* from different models vary in the range of 1.6–4.4, depending on the nature of cross-link interactions and network topology and dimensionality [25, 59–63]. In our experiments, the measured exponent *β* = 0.18±0.11 for PrLD-SAM is much smaller than previous theoretical estimates. A possible reason for this discrepancy is that the control parameter *ϕ* used in the experiment differs from those used in theories. In percolation theories, the volume fraction *ϕ* (or probability *p*) falls in the range between 0 and 1. When *ϕ* = 0 (or *p* = 0), no bond is formed and the system remains to be completely floppy, similar to ideal gases or liquids. When *ϕ* = 1 (or *p* = 1), all elements in the system are connected and it corresponds to an infinitely-rigid solid. The molar fraction *ϕ* used in this experiment, however, is the fraction of WT in binary MH-WT condensates, and therefore only covers a limited range of theoretical *ϕ*. Moreover, the protein network is very different from the lattice models of percolation with permanent crosslinks. The bonds in protein condensates are dynamic and can unbind over longer timescales. This dynamic nature of protein cross-links requires new theoretical calculation of the critical exponent *β*.

Our analysis of the critical behavior of PrLD-SAM condensates carries important biological implications, as it sheds new light on how protein condensates maintain their mechanical homeostasis, a balance between structural robustness and flexibility. In particular, the protein network formed by wild-type SAM resides near the critical threshold, with its apparent modulus *E* positioned on the rising shoulder of the transition curve. This renders the condensate in a distinctive viscoelastic regime where elasticity and viscosity contribute nearly equally. At long times, the measured relaxation modulus follows a power-law form, *E*(*t*) ≃ *E*_2_(*t/τ*_2_)^*−α*^, with a power law exponent *α* ≃ 0.5 (see Eq. (2)) [57]. The corresponding storage and loss moduli go as *G*^*′*^(*f*) ∼ *G*^*′′*^(*f*) ∼ (2*πfτ*_2_)^*α*^ and their amplitude ratio is governed by the phase angle,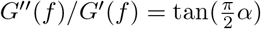. For *α* = 1*/*2, the ratio equals 1, marking a crossover between solid-like behavior (elastic dominance, *G*^*′′*^*/G*^*′*^ < 1 when *α* < 1*/*2) and liquid-like behavior (viscous dominance, *G*^*′′*^*/G*^*′*^ > 1 when *α* > 1*/*2). Thus, the condensate network in WT indeed resides at this viscoelastic crossover point.

Biological condensates require distinct material properties to support their multifaceted physiological functions. On one hand, an adequate condensate elasticity is essential for maintaining the structural integrity and physiological functions of protein condensates. In living cells, network structures, such as the cytoskeleton and endoplasmic reticulum (ER), play essential roles in preserving cellular integrity, mechanical stability, and homeostasis [64–67]. Analogously, protein condensates, particularly those stable assemblies such as synaptic condensates that can last for decades, rely on a percolated network with sufficient mechanical stiffness to resist external perturbations and prevent entropic mixing with their environment [13, 19, 68]. A purely fluid-like state, in which the molecules undergo random Brownian motion, would severely affect condensate structure and functions.

This network-based structural integrity is particularly important for the PSD condensate, which helps localize synaptic receptors and support synaptic signaling [14, 19, 69, 70]. Disruption or loss of this percolated network jeopardizes the normal functions of the PSD. Pathological mutations observed in human patients weaken SAM-SAM interactions, causing the network to collapse in PrLD-SAM condensates (see Fig. 7). Our recent work further showed that Shank3 mutations that impair Shank oligomerization soften the PSD condensate by destabilizing its percolated network, resulting in impaired synaptic transmission and autism-like behavior in mouse model [19].

On the other hand, functional condensates require a sufficient degree of internal fluidity and flexibility for normal physiological functions, such as those in actively responding to signaling cues [16, 71]. An overly rigid network or structure would severely limit molecular mobility, thereby suppressing condensate activities and disrupting their normal functions, such as DNA damage repair [71], condensate-mediated enzyme catalysis [72], and cellular stress response [73]. The aging-related transition from a fluidic/dynamic state to a solid-like irreversible state is also a notorious example. The loss of fluidity in the fibril tangles in the nervous system can severely impair cellular functions and disrupt biochemical regulations, which potentially leads to pathological changes and the onset of neurodegenerative diseases [1, 15, 16]. Our findings thus suggest that protein condensates operate at a narrow range of optimal mechanical states near the rigidity percolation threshold, where a cohesive yet compliant network provides structural robustness while maintaining flexibility essential to condensate function.

Functional condensates staying in a near-critical state is an intriguing phenomenon in line with the hypothesis of self-organized criticality, where living systems spontaneously evolve toward criticality without external finetuning [74–77]. Other examples include neuronal signal propagation [78], gene regulatory network [79] and collective motion of bird flocks [80]. Far from the critical point, molecular interactions in condensates are predominantly local and short-ranged. In this regime, the central limit theorem applies: local random fluctuations average out over large scales, rendering macroscopic material properties, such as the Young’s modulus, effectively decoupled from specific microscopic (molecular) details and determined instead by statistical averages over vast numbers of degrees of freedom. Near the critical point, however, the correlation length diverges (theoretically approaching infinity as the control parameter reaches criticality), leading to long-range correlations that invalidate the central limit theorem [81, 82]. Microscopic details are no longer averaged out but become directly amplified and coupled to macroscopic properties through emergence.

Operating within this critical regime grants systems exquisite sensitivity: small molecular-scale perturbations, such as biochemical signaling or mutations, can propagate via amplified correlations to trigger system-wide structural or mechanical transitions [77, 79]. This enhanced responsiveness, inherent to criticality, underpins the fundamental homeostasis, adaptability and plasticity of living systems. Evolution may have harnessed these critical states to foster biological complexity, as single-point mutations are capable of generating disproportionately large phenotypic effects [83, 84]. Conversely, deviations from criticality, such as the disease-associated mutant LW, which falls below the percolation threshold (see Fig. 7), can lead to the collapse of condensate structures and subsequent pathological changes.

The critical behavior observed in PrLD-SAM condensates, combined with their capacity to maintain balanced viscoelasticity near the percolation point, underscores the remarkable adaptability inherent in protein condensates. These principles likely extend to the broader cellular context, providing a unifying framework for understanding how biomolecular assemblies achieve the dual requirements of robustness and flexibility within complex biological environments.

## Acknowledgments

This work was supported, in part, by Shenzhen Medical Research Fund under grant no. E250200113 (M.Z.) and RGC of Hong Kong SAR under grant nos. 16302025 (P.T.), 16300224 (P.T.), and C6041-24G (P.T.).

## Author contributions

Z.L., M.Z. and P.T. conceived ideas and designed research; Z.L. conducted AFM experiments and AFM data analysis; B.J., Y.X., and Z.L. conducted fluorescent and biochemical experiments; B.J., Y.X. and Z.S. performed SMT experiments; all authors analyzed and interpreted data; Z.L. and P.T. drafted the paper with inputs from all other authors; M.Z. and P.T. coordinated the project.

## Data availability

Source data that support the findings of this study (such as figure source data) are provided with this paper in the Source Data file. Unless otherwise stated, all the data supporting the results of this study can be found in the article, supplementary, and source data files. The raw data is available from the corresponding authors upon request. Source data is provided with this paper.

## Notes

### Competing Interest Statement

The authors have declared no competing interest.

